# Parkinson’s disease-associated alterations of the gut microbiome can invoke disease-relevant metabolic changes

**DOI:** 10.1101/691030

**Authors:** Federico Baldini, Johannes Hertel, Estelle Sandt, Cyrille C. Thinnes, Lorieza Neuberger-Castillo, Lukas Pavelka, Fay Betsou, Rejko Krüger, Ines Thiele, on behalf of the NCER-PD Consortium

## Abstract

Parkinson’s disease (PD) is a systemic disease clinically defined by the degeneration of dopaminergic neurons in the brain. While alterations in the gut microbiome composition have been reported in PD, their functional consequences remain unclear. Herein, we first analysed the gut microbiome of patients and healthy controls by 16S rRNA gene sequencing of stool samples from the Luxembourg Parkinson’s study (n=147 typical PD cases, n=162 controls). All individuals underwent detailed clinical assessment, including neurological examinations and neuropsychological tests followed by self-reporting questionnaires. Second, we predicted the potential secretion for 129 microbial metabolites through personalised metabolic modelling using the microbiome data and genome-scale metabolic reconstructions of human gut microbes. Our key results include: 1. eight genera and nine species changed significantly in their relative abundances between PD patients and healthy controls. 2. PD-associated microbial patterns statistically depended on sex, age, BMI, and constipation. The relative abundances of *Bilophila* and *Paraprevotella* were significantly associated with the Hoehn and Yahr staging after controlling for the disease duration. In contrast, dopaminergic medication had no detectable effect on the PD microbiome composition. 3. Personalised metabolic modelling of the gut microbiomes revealed PD-associated metabolic patterns in secretion potential of nine microbial metabolites in PD, including increased methionine and cysteinylglycine. The microbial pantothenic acid production potential was linked to the presence of specific non-motor symptoms and attributed to individual bacteria, such as *Akkermansia muciniphila* and *Bilophila wardswarthia*. Our results suggest that PD-associated alterations of gut microbiome could translate into functional differences affecting host metabolism and disease phenotype.

## INTRODUCTION

Parkinson’s Disease (PD) is a complex multifactorial disease, with both genetic and environmental factors contributing to the evolution and progression of the disease (Kalia et al. 2015). While several studies have elucidated the role of genetic factors in the pathogenesis of the disease (Kitada et al. 1998; Bonifati et al. 2003; Paisan-Ruiz et al. 2004; Di Fonzo et al. 2009), the role and the contribution of various environmental and lifestyle factors are still not completely understood (Gatto et al. 2010). Importantly, about 60% of the PD patients suffer from constipation (Fasano et al. 2015), which can start up to 20 years before the diagnosis and is one of the prodromal syndromes (Savica et al. 2009; Cersosimo et al. 2013).

The human being is considered to be a superorganism recognising a complex interplay between the host and microbes (Sleator 2010). For instance, the human gut microbiome has been shown to complement the host with essential functions (trophic, metabolic, protective) and to influence the host’s central nervous system (CNS) via the gut-brain axis through the modulation of neural pathways and GABAergic and serotoninergic signalling systems (Carabotti et al. 2015).

Recent studies have reported an altered gut composition in PD (Hasegawa et al. 2015; Keshavarzian et al. 2015; Scheperjans et al. 2015; Bedarf et al. 2017; Hill-Burns et al. 2017; Hopfner et al. 2017; Petrov et al. 2017; Heintz-Buschart et al. 2018; Barichella et al. 2019). One of these studies has been conducted using samples from recently diagnosed, drug-naive patients (Bedarf et al. 2017). These studies have demonstrated that PD patients have an altered microbiome composition, compared to age-matched controls. However, the functional implications of the altered microbiome remain to be elucidated, e.g., using animal models (Sampson et al. 2016). A complementary approach is computational modelling, or constraint-based reconstruction and analyses (COBRA) (Orth et al. 2010), of microbiome-level metabolism. In this approach, metabolic reconstructions for hundreds of gut microbes (Magnusdottir et al. 2017) are combined based on microbiome data (Baldini et al. 2018; Heirendt et al. 2019)). Flux balance analysis (FBA) (Orth et al. 2010) is then used to compute, e.g., possible metabolite uptake or secretion flux rates of each microbiome model (microbiome metabolic profile) (Heinken et al. 2019) or to study of microbial metabolic interactions (cross-feedings) (Klitgord and Segre 2010; Heinken and Thiele 2015). This approach has been applied to various microbiome data sets to gain functional insights (Thiele et al. 2018; Heinken et al. 2019; Hertel et al. in revision), including for PD where we propose that microbial sulphur metabolism could contribute to changes in the blood metabolome of PD patients (Hertel et al. in revision).

In the present study, we aim at investigating microbial changes associated with PD while focusing on possible covariates influencing microbial composition and at proposing functional, i.e., metabolic, consequences arising from the microbiome changes. First, we analysed the faecal microbial composition of PD patients and controls from the Luxembourg Parkinson’s study (Hipp et al. 2018) (Figure 1). Second, based on the observed significant differences in the composition of microbial communities between PD patients and controls, we created and interrogated personalised computational models representing the metabolism of each individual’s microbial community. We demonstrate that the combined microbial composition and functional metabolite analysis provides novel hypotheses on microbial changes associated with PD and disease severity, enabling future mechanism-based experiments.

**Figure 1:**
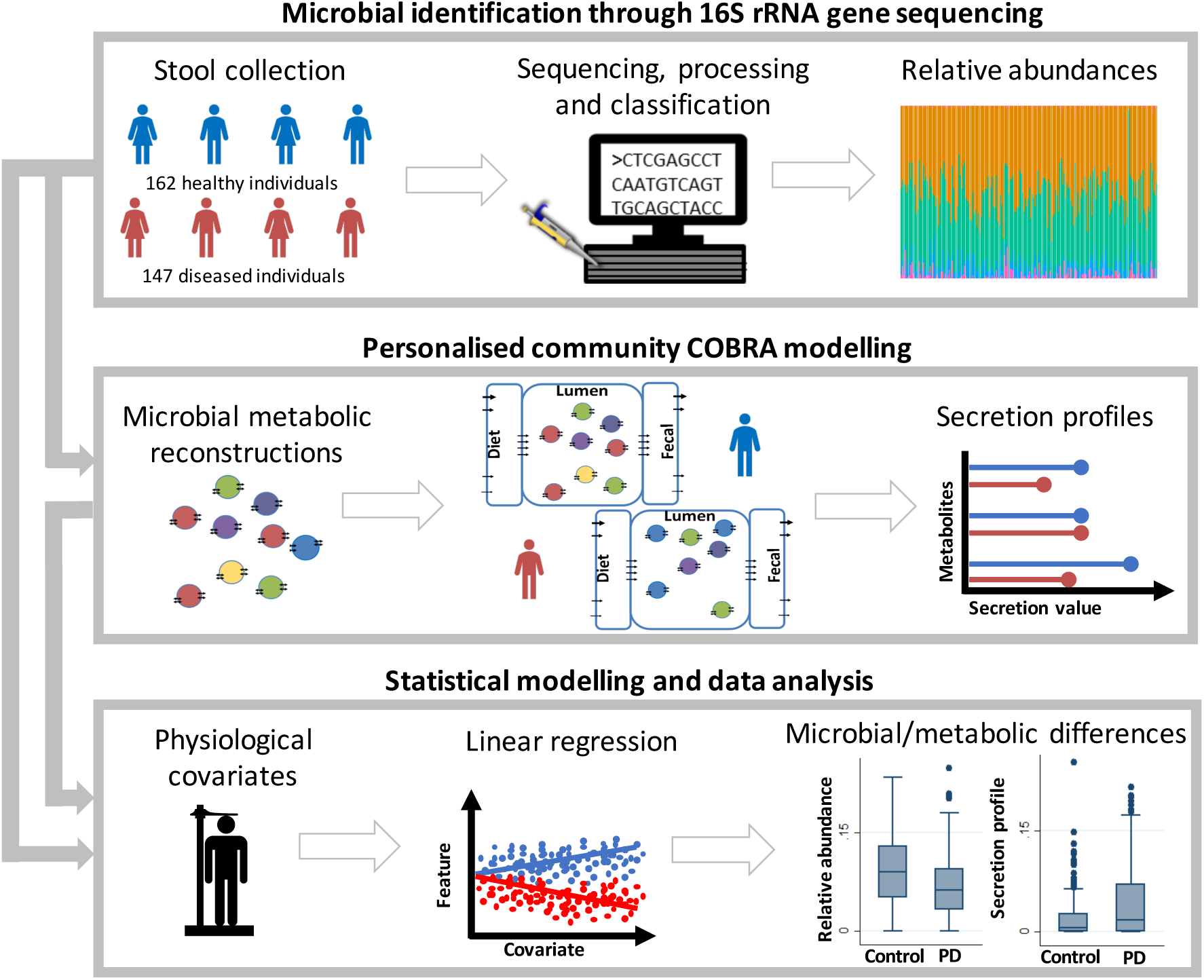
Overview of the study approach and the key methods used. Relative abundances were derived from 16S rRNA gene sequences (Methods: Analysis of the microbial composition with 16S rRNA gene sequencing) and used as input for the personalised community modelling to simulate metabolites secretion profiles. Relative abundances and secretion profiles were statistically analysed to identify microbial or metabolic differences between PD patients and controls.

## RESULTS

The Luxembourg Parkinson’s Disease study includes patients with typical PD and atypical parkinsonism, as well as matched healthy control subjects from Luxembourg and its neighbouring regions from a broad age-range (Hipp et al. 2018). For the present study, we focused on typical PD patients and healthy controls over the age of 50 (Table 1, Methods). Stool samples were analysed for 147 PD patients and 162 controls using 16S rRNA gene sequences (Methods: Analysis of the microbial composition with 16S rRNA gene sequencing).

**Table 1:**
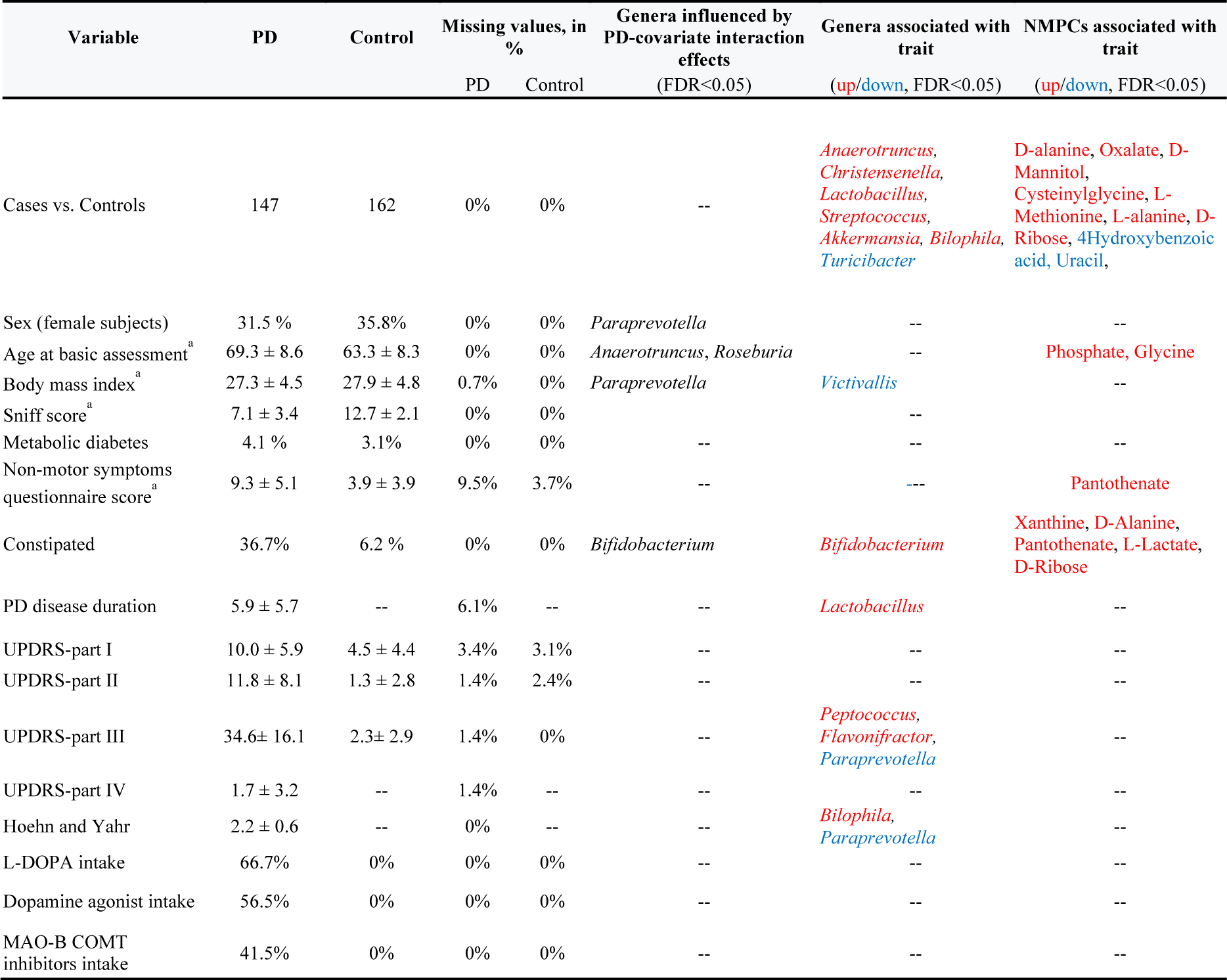
Descriptive statistics of the analyses sample from the Luxembourg Parkinson’s Disease study and overview over associations. A red label means increased in PD, blue decreased in PD, while -- “nothing to report”. PD disease duration refers to time since diagnosis at the date of stool sampling. UPDRS=Unified Parkinson Rating Scale, L-DOPA=levodopa, MAO-B=monoaminooxidase B, COMT=Catecholamine-Methyl-Transferase, NMPC=Net maximal production capability.

### Species and genus level changes in PD microbiomes

We investigated disease-associated microbial changes at the species level. We found that the mean species diversity (i.e., the alpha-diversity) did not significantly differ between PD cases and controls (b=-0.04351, 95%-CI:(−.107;0.177), p=0.177), in agreement with earlier studies (Scheperjans et al. 2015; Bedarf et al. 2017) (Hopfner et al. 2017), but in disagreement with two other studies (Keshavarzian et al. 2015; Heintz-Buschart et al. 2018). However, seven species were significantly altered in PD (FDR<0.05, Figure 2). Note that when comparing results between different taxonomic levels, changes observed for *Ruminococcus* and *Roseburia* species were not significant on the genus level but only on the species level, highlighting the importance of species-level resolution. The highest effect size was associated with *Akkermansia muciniphila* (Odds ratio (OR)=1.80, 95%-CI=(1.29, 2.51), p=6.02e-04, FDR<0.05; Supplementary Table 1) in agreement with the previously reported higher abundance of *A. muciniphila* in PD patients (Bedarf et al. 2017; Heintz-Buschart et al. 2018)). Subsequently, we examined possible differences at the genus level by performing semiparametric fractional regressions while adjusting for age, sex, the body mass index (BMI), batch, and total read counts. We identified eight genera to be significantly increased in PD (FDR<0.05; Figure 3A, Table 1), with *Lactobacillus* showing the highest effect size (Odds ratio (OR)=5.75, 95%-CI=(2.29, 14.45), p=1.96e-04, FDR<0.05; Supplementary Table 2). In contrast, the genera *Turicibacter* decreased significantly in PD cases (FDR<0.05). To summarise, significant changes could be observed on the species and genus level.

**Figure 2:**
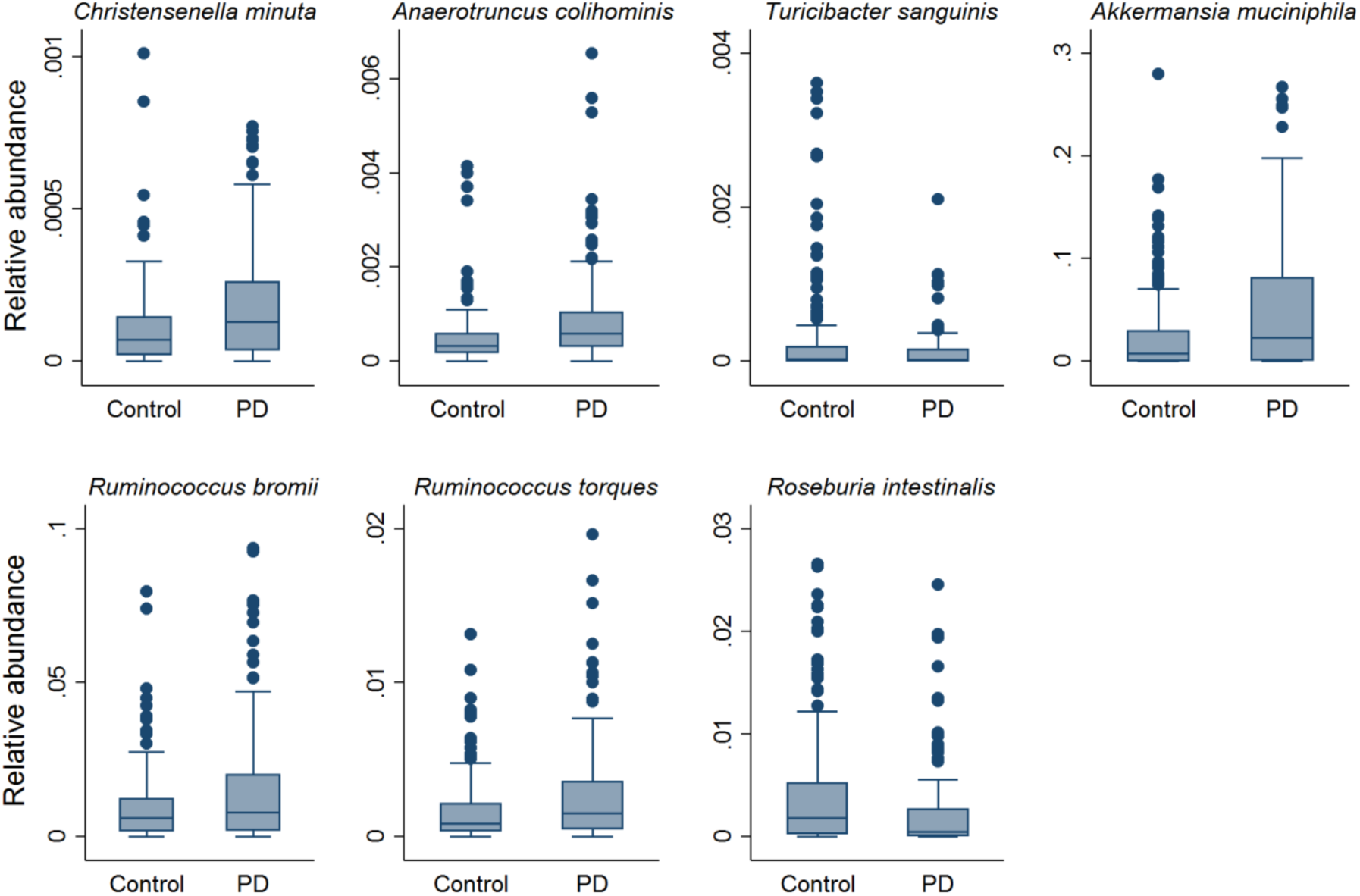
Boxplots of seven significantly changed species in PD versus controls. (FDR<0.05). Significance levels were determined using multivariable semi-parametrical fractional regressions with the group variable (PD vs. control) as predictor of interest, including age, gender, BMI, and technical variables (i.e., total read-counts and batch effect) as covariates. FDR=false discovery rate.

**Figure 3:**
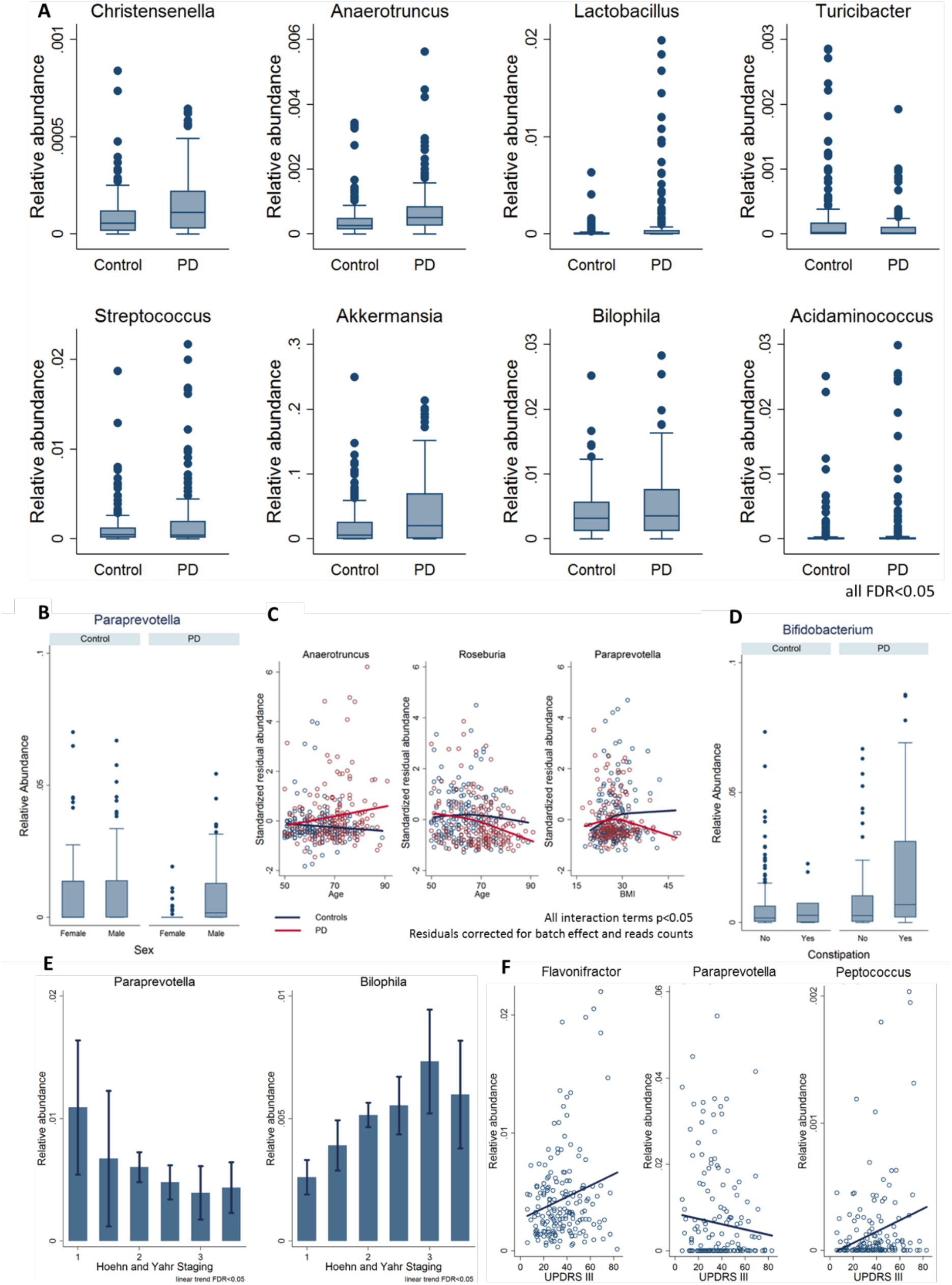
Genus alterations in PDs due to interactions with basic covariates. Relative abundance is given on a logarithmic scale. **A.** Boxplots of the seven significant species (FDR<0.05). **B.** Female PD patients have a reduced abundance of *Paraprevotella* (FDR<0.05). **C.** Genus abundance age and BMI dependencies of *Anaerotruncus, Roseburia*, and *Paraprevotella* (global test on all interaction terms, FDR<0.05). For graphical assessment of the interaction terms the z-transformed residual abundances are displayed after correction for technical covariates (batch and read counts). **D.** The genus relative abundance of Bifidobacterium was increased in patients reporting to be constipated (FDR<0.05). **E.** Genus association with disease staging showed a decrease of relative abundance of *Paraprevotella* and an increase of *Bilophila* genus over increasing Hoehn and Yahr scale values (FDR<0.05). **F.** An increased score in motor symptoms (UPDRS III) was associated with an increased trend in abundances of *Flavonifractor* and *Peptococcus* and a decreased trend in *Paraprevotella* abundance (FDR<0.05). UPDRS=Unified Parkinson Rating Scale, BMI=body mass index, FDR=false discovery rate.

### PD modifies the effects of basic covariates on the microbiome

Furthermore, we investigated whether the genus level alterations in PD were affected by basic confounding factors. This interaction analyses uncovered rich effect modifications, revealing that microbiome changes in PD have to be considered in the context of age, BMI, and gender. Our analyses demonstrate that the effects of PD are not homogeneous among important sub-groups of patients. For example, *Paraprevotella* was exclusively reduced in female patients but not in female controls (Figure 3B), highlighting gender-dependent alterations of microbial communities in PD. In addition, the effects of BMI and age were modified in PD cases. The PD cases had increased *Anaerotruncus* abundance with age, while non-linear, overall decreasing abundances of *Roseburia* and *Paraprevotella* were observed with age and BMI, respectively (Figure 3C). Taken together, these analyses suggest that microbial abundances are shifted in PD cases and that also the effects of important covariates were altered in PD, reflecting the systemic and complex nature of PD.

### Microbial abundances, medication intake, and constipation in PD

The Luxembourg Parkinson’s study enrols patients of all stages of PD. Therefore, the patients have considerable inter-individual variance in PD-related features, such as constipation and intake of medication (Table 1). We analysed whether these features had an impact the microbiome composition in PD. In our data, we could not find any evidence for an effect of the three medication types on the microbiome, i.e., levodopa, COMT inhibitors, or MAO-B inhibitors, when correcting for multiple testing (Supplementary Table 2). In contrast, constipation, a prevalent non-motor symptom in PD patients (Lesser 2002), was associated with an increased abundance of *Bifidobacterium*, with a clear effect in constipated PD cases (Figure 3D). However, since there were only ten constipated controls (Table 1), these results must be confirmed in larger cohorts.

### Genus association with the disease severity

We next investigated whether the stage of the disease, i.e., defined by Hoehn and Yahr staging, NMS, and UPDRS (Unified Parkinson Rating Scale) scores, and its subscales, was associated with altered genus abundance. For the Hoehn and Yahr staging, *Paraprevotella* showed a negative association and *Bilophila* showed a positive association, both of which were significant after multiple testing (Figure 3E). For the UPDRS III subscale score (i.e., motor symptoms, Table 1), three genera, being *Peptococcus, Flavonifractor*, and *Paraprevotella*, survived correction for multiple testing (Figure 3F). In contrast, the other UPDRS subscales and the NMS were not significantly associated with microbial changes, after correction for multiple testing. Note that these analyses were performed while adjusting for disease duration. When analysing the association pattern of disease duration, we found *Lactobacillus* positively correlated with the disease duration (FDR<0.05, Supplementary Figure S1). In conclusion, our data suggest that the microbial composition may be utilised as a correlate of disease severity.

### Metabolic modelling reveals distinct metabolic secretion capabilities of PD microbiomes

To obtain insight into the possible functional consequence of observed microbiome changes in PD, we used metabolic modelling (cf. Methods). Briefly, we mapped each of the 309 microbiome samples on the generic microbial community model consisting of 819 gut microbial reconstructions (Magnusdottir et al. 2017; Heinken et al. 2019) (cf. Supplementary Material) to derived personalised microbiome models (Baldini et al. 2018). We then computed a net maximal production capability (NMPC) for 129 different metabolites that could be secreted by each microbial community model (cf. Methods), providing thereby a characterisation of the differential microbial metabolic capabilities in PDs and controls. The secretion of nine metabolites had differential NMPCs in PD (Figure 4A, all FDR<0.05) as determined by multivariable regressions adjusting for age, sex, BMI, and technical covariates. Moreover, although less dominant in comparison to the abundance data, PD-covariate interactions were also prevalent, with the uracil secretion potential showing a sex-specific effect and cysteine-glycine showing a BMI-dependent PD-effect (Figure 4B, 4C). In subsequent analyses, we tested for associations of the NMPCs with constipation, medication, disease duration, Hoehn-Yahr staging, NMS, and UPDRS III scores, complementing thereby the analyses on the abundance level. Notably, we found xanthine, D-alanine, L-lactic acid, D-ribose, and pantothenic acid positively associated with constipation (Figure 4B), while no NMPC was associated with medication or with disease duration. However, the pantothenic acid secretion potential was positively associated with higher NMS scores, interestingly both in PD and in controls (Figure 4D), while no NMPC survived correction for multiple testing regarding associations with the UPDRS III score and Hoehn-Yahr staging. To conclude, these results suggest that the altered microbial composition in PD could result in broad changes in metabolic capabilities, which manifested themselves additionally in non-motor symptoms and constipation.

**Figure 4:**
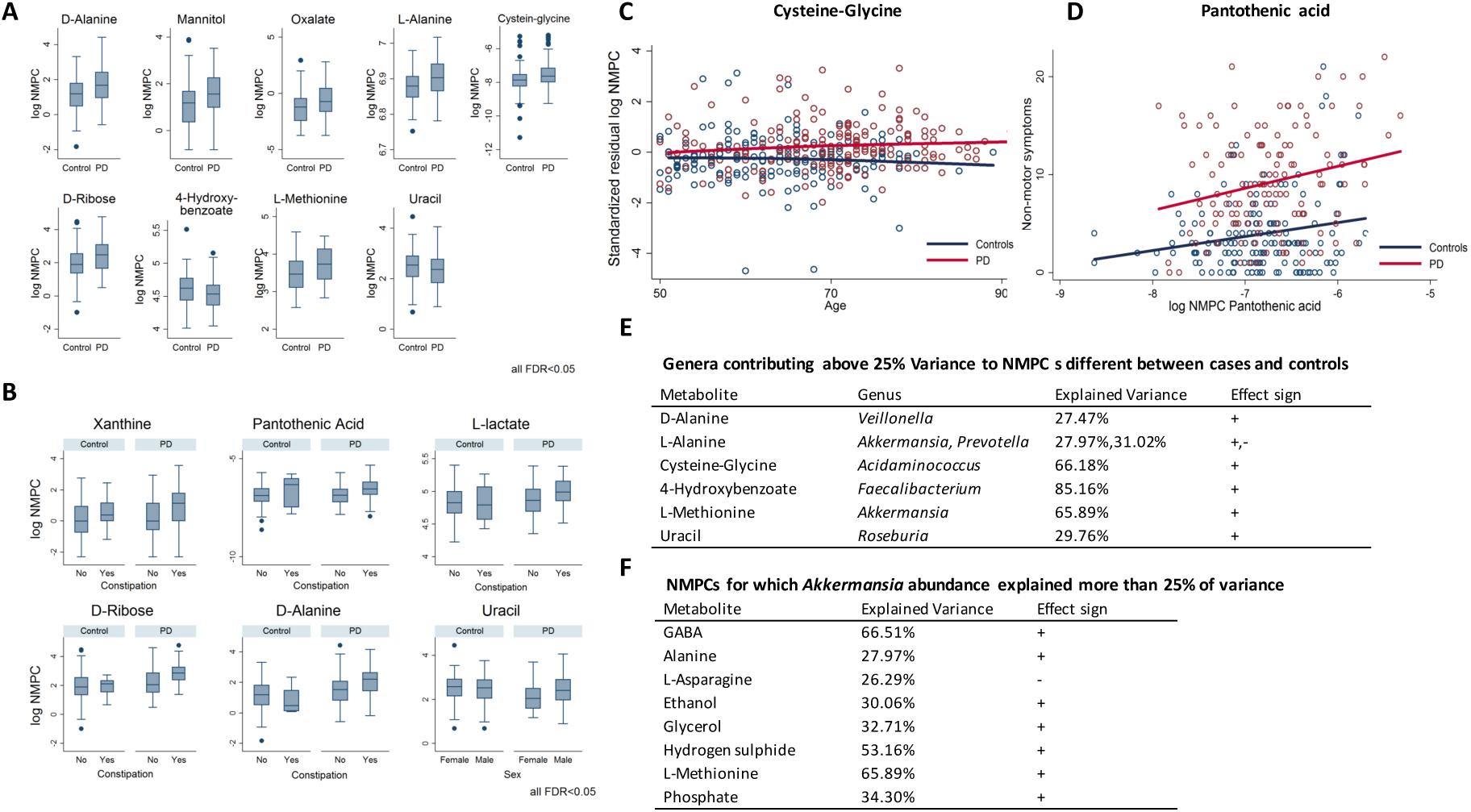
Result of analysing secretion profiles of microbial communities. **A.** Box plots for NMPCs differential between cases and controls with FDR<0.05. **B.** NMPCs with sex-specific PD signature or constipation effects (all FDR<0.05). **C.** Differential age trajectory between cases and controls for cysteine-glycine (p<0.05). **D.** Association of pantothenic acid with non-motor symptoms. **E.** Genera contributing more than 25% to NMPCs different between cases and controls. **F.** *Akkermansia* contribution to community production of 12 metabolites expressed as a percentage of total production for each compound. Metabolites highlighted in red were significantly increased in PD (FDR<0.05). NMPC=net maximal production capacity, GABA=gamma-aminobutyrate, H2S=hydrogen sulphide, FDR=false discovery rate. Effect sign “–”: negative correlation. Effect sign “+”: positive correlation.

### PD specific secretion profiles were altered due to changed community structure and species abundances

Next, we analysed which microbes contributed to the differential secretion profiles by correlating the NMPCs to the abundance data (Figure 4E/F, Supplementary Table 3). Six metabolite NMPCs had strong contribution or where even dominated by single genera (Figure 4D), while for the other four NMPCs no single dominant genus could be identified. We then computed the contribution value of each genus to the production of each secreted metabolite (NMPC). From the aforementioned genera, which were associated on genus or species level with PD, only *Akkermansia, Acidaminococcus*, and *Roseburia* had substantial metabolic contributions (over 25%). *Acidaminococcus* was responsible for 64% of the variance in cysteine-glycine production and *Roseburia* for 30% of the variance in uracil production potential. *Akkermansia* impacted the secretion profiles the most and substantially contributed to the metabolism of nine metabolites (Figure 4F), including the neurotransmitter gamma-aminobutyric acid (GABA) and two sulphur species, being hydrogen sulphide and methionine. GABA was also significantly altered between PD and controls on a nominal level missing FDR corrected significance narrowly (b=0.18, 95%-CI:(0.06;0.30), p=0.003, FDR=0.0501). These analyses demonstrate the added value of metabolic modelling to investigate altered metabolic functions from the whole microbial composition.

## DISCUSSION

In this study, we aimed to elucidate compositional and functional changes in the faecal microbiome of PD patients. Therefore, we analysed 16S rRNA data from a cohort of typical PD patients (n=147) and controls (n=162), and performed personalised microbial computational modelling. We identified i) eight genera and nine species that changed significantly in their relative abundances between PD patients and healthy controls. ii) PD-associated microbial patterns that were dependent on sex, age, BMI, constipation, and iii) in PD patients altered secretion potentials, particularly in sulphur metabolism, using metabolic modelling of microbial communities. Overall, our work demonstrated compositional and functional differences in the gut microbial communities of Parkinson’s disease patients providing novel experimentally testable hypothesis related to PD pathogenesis.

The microbial compositional analyses of our cohort identified significantly different microbial abundance distributions between PD patients and healthy controls (Table 1). Up to date, 13 studies have described altered colonic microbial compositions associated with PD and an overall picture starts to arise (Figure 5). For instance, the microbial the families of Verrucomicrobiaceae and Lactobacillaceae have been consistently found to have an increased abundance in PD (Figure 5). In accordance, our study also reports increased abundance in PD of Akkermansia, Christensenella, and Lactobacillus. Similarly, Bifidobacteria has also been repeatedly associated with PD (Figure 5) but in our study, we could show that the Bifidobacteria association dependent on constipation (Figure 3) highlighting the need for incorporating disease-specific phenotypes as covariates into the statistical design.

**Figure 5:**
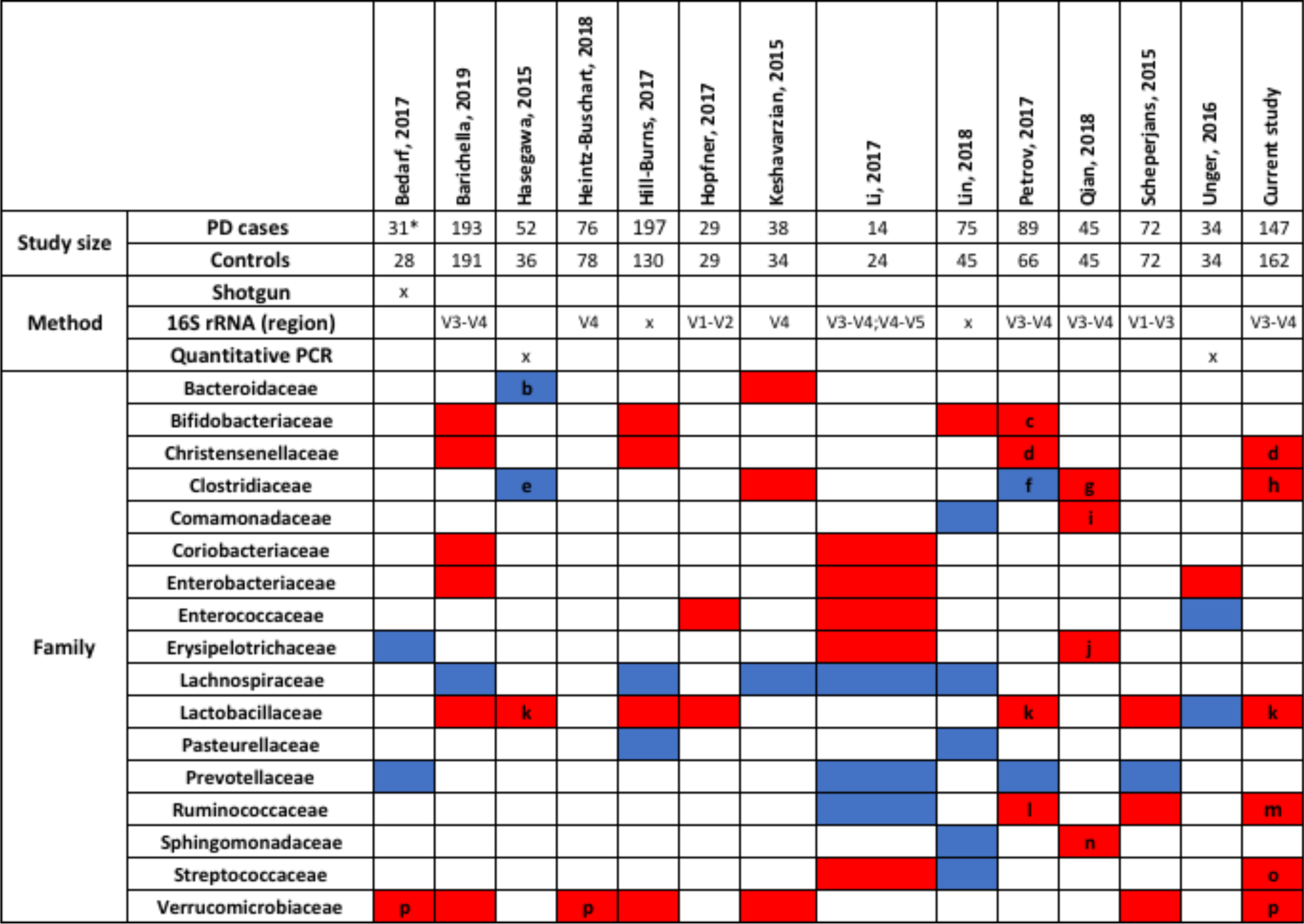
Reported microbial changes at the family level associated with PD in different studies. Only those bacterial families are shown, for which significant associations with species or genera have been reported in at least two studies comparing stool samples from patients and controls. Red - increased in PD, Blue - decreased in PD. a: *Actinomycetales*, b: *Bacteroides fragilis*, c: *Bifidobacterium*, d: *Christensenella*, e: *Clostridium coccoides*/*leptum*, f: *Faecalibacterium, Dorea*, g: *Clostridium* IV/XVIII, *Butyricicoccus, Anaerotruncus*, h: *Anaerotruncus*, i: *Aquabacterium*, j: *Holdemania*, k: *Lactobacillus*, l: *Oscillospira*, m: *Ruminococcus romii, Ruminococcus torques*, n: *Sphingomonas*, o: *Streptococcus*, p: *Akkermansia*. * Drug-naive, de novo PD patients only. Based on (Barichella et al. 2019).

At the same time, inconsistencies between the studies remain and they may be due to differences in study design, inclusion criteria, faecal sampling, RNA extraction protocols, and metagenomic and statistical methods. For instance, we used a relatively large, PD cohort while Bedarf and colleagues (Bedarf et al. 2017) studied a small cohort of drug-naïve, male PD patients and male controls (Figure 5). Three studies included individuals of Chinese descent (Li et al. 2017; Lin et al. 2018; Qian et al. 2018) while the other studies focused on Caucasian individuals. It has been shown that microbial composition is associated with ethnic background, geography, and dietary habits (Turnbaugh et al. 2008; De Filippo et al. 2010; Yatsunenko et al. 2012), which may explain some of the discrepancies. The differences between the studies hence highlight the importance of performing meta-analysis to identify global microbial signatures, as it has been done for, e.g., colorectal cancer (Wirbel et al. 2019). Such meta-analysis may also permit to investigate subgroups of PD, as the number of cases and controls would be substantially increased and thus provide higher statistical power. For instance, we observed various effect modulators that were not reported before in humans (Table 1), such as *Paraprevotella* abundance reduction being specific to women. This result is apparently in contradiction with findings from Bedarf and colleagues (Bedarf et al. 2017) who reported decreased levels of Prevotellaceae in a cohort of only male PD patients. However, once again, differences might be explained by different inclusion criteria, methodologies, and related possible sex-specific effects. Interestingly, a recent study reported a higher abundance of *Paraprevotella* in male mice compared to female mice (Huang et al. 2018). Despite the lack of extensive studies on gender-specific differences in microbiome composition, we suggest that machine learning procedures on microbiome data should be performed in a sex-stratified manner. Larger cohorts, e.g., through meta-analysis of published cohorts would allow the identification of generalizable microbial differences in PD patients and also, specific microbial changes associated with certain traits and physiological characteristics, as suggested by our data.

We could not detect an effect of the dopaminergic, PD specific medication on the microbiome composition, after correction for multiple testing. Also the fact that key findings from the study of Bedarf and colleagues were reproduced in other cohorts of PD patients under medication, including ours, support that notion. Nonetheless, in previous studies, Dorea and Phascolarctobacterium genera have been negatively associated with levodopa equivalent doses (Qian et al. 2018) and members of the family of *Bacillaceae* have been correlated with levodopa treatment (Heintz-Buschart et al. 2018). Consequently, it cannot be excluded that medication is associated with microbial changes, albeit the association may be weaker than the effects of other covariates. As PD drugs are often taken in combinations, it would require a larger sample size than used in our study to permit the investigation of all possible drug combinations. The lack of clear association is somewhat expected as levodopa is absorbed in the upper part of the small intestine (Streubel et al. 2006) and thus small intestinal rather than large intestinal microbes may play a more prominent role in levodopa bioavailability. Consistently, a recent study showed that bacterial tyrosine decarboxylases restrict the bioavailability of levodopa (van Kessel et al. 2019). Interestingly, 193/818 reconstructed microbes (Magnusdottir et al. 2017), commonly found in the human gut, carry genes encoding for proteins that convert levodopa into dopamine (Noronha et al. 2019). Levodopa is always given with decarboxylase inhibitors, such as carbidopa or benserazide, targeting the human decarboxylases, but it cannot be excluded that they also act on the microbial counterpart. However, Van Kessel et al. have shown that carbidopa as well as benserazide is only a weak inhibitor of the microbial tyrosine decarboxylase (van Kessel et al. 2019).

We identified a positive association of *Bilophila* abundance with the Hoehn and Yahr staging, which captures motor impairment and disability independent of disease duration. Indeed, the abundance of *Bilophila* was not associate with disease duration, indicating mainly dependency on the progression of symptoms. This finding is consistent with experimental mice studies demonstrating the pro-inflammatory effect of *Bilophila* overgrowth (Devkota et al. 2012; Natividad et al. 2018). Notably, the Hoehn and Yahr staging was also positively associated on a nominal level with the predicted pyruvate secretion profile (Supplementary File 4), which was accordingly significantly increased in PD patients on a nominal level alongside with L- and D-alanine. *Bilophila* has the rare capability to use taurine, an inhibitory neurotransmitter with neuroprotective effects (Saransaari and Oja 2007; Wu et al. 2009), as an energy source (Laue and Cook 2000). This pathway is initiated by the taurine: pyruvate aminotransferase (Laue and Cook 2000), converting pyruvate and taurine into L-alanine and sulfoacetaldehyde. The only microbe of the 818 species in our AGORA collection encoding the corresponding gene was *Bilophila*, which was significantly increased (FDR<0.05) and hence, the corresponding reaction (VMH ID: TAURPYRAT) was increased in abundance in PD microbiomes as well. In a previous study (Hertel et al. in revision), we have shown that blood taurine conjugated bile acids were positively associated with motor symptoms. *Bilophila* may be a marker of disease progression in PD, and it could modulate human sulphur metabolism through its taurine degradation capabilities. Alterations in sulphur metabolism have been already described when using computational modelling of microbiomes from a cohort of early diagnosed and levodopa naive PD patients (Bedarf et al. 2017; Hertel et al. in revision) as well as an increased concentration of methionine and derived metabolites in blood samples (Hertel et al. in revision). Furthermore, we and others have reported alterations in bile acids and taurine-conjugated bile acids in PD patients (Graham et al. 2018; Hertel et al. in revision). Our present study suggests again a key role of *Bilophila* in host-microbiome sulphur co-metabolism, which may link with bile acid metabolism.

Interestingly, an increased abundance of *B. wadsworthia* has been linked to constipation (Vandeputte et al. 2017). *B. wadsworthia* is the only microbe in the AGORA collection capable of the metabolic reaction converting pyruvate and taurine to L-alanine and sulphoacetaldehyde (VMH ID: TAURPYRAT). Therefore, an increased production of L-alanine might be due to the increased *B. wadsworthia* abundance. This resulting higher production rate of L-alanine could then lead to an increased conversion into D-alanine via the alanine racemase (VMH ID: ALAR), which was present in 808/818 gut microbes in the AGORA collection. Accordingly, D-alanine was one of the three metabolite secretion profiles increased in constipated PD patients (Figure 4E). This hypothesis of *B. wadsworthia* playing a role in constipation of PD patients would need to be experimentally validated, especially since we could not find statistically significant changes in the association between the abundance of *B. wadsworthia* and constipated individuals. In contrast, we found an increase in *Bifidobacteria* abundance in constipated individuals and particularly in constipated PD patients. This result disagreed with an earlier study on individuals with chronic constipation, which reported a decrease in *Bifidobacteria* abundance (Khalif et al. 2005). Overall, the available data suggest that complex alterations in microbial composition are associated with constipation but may differ between diseases.

The mucin degrading microbe, *A. muciniphila*, represents about 1-4% of the faecal microbiome in humans (Naito et al. 2018). Numerous diseases have been associated with a decrease in *A. muciniphila* abundance (Schneeberger et al. 2015; Grander et al. 2018), while an increase has been consistently reported in PD patients (Figure 5). The *A. muciniphila* abundance had the largest contribution to the significantly altered metabolite secretion profiles (Figure 4E), including the neurotransmitter gamma-aminobutyric acid (GABA). While its predicted secretion potential was only nominally increased in PD patients the present study, higher GABA secretions rates have also been predicted based on microbiome data from early stage levodopa naive PD patients (Hertel et al. in revision). Importantly, GABA receptors have been found in the enteric nervous system, gut muscle, gut epithelial layers, and endocrine-like cells (Hyland and Cryan 2010) and its gut receptors are thought to be related to gastric motility (peristalsis), gastric emptying, and acid secretion (Hyland and Cryan 2010). Experiments with the GABAb agonist baclofen have shown that GABAb receptors can reduce gastric mobility in the colon of rabbits (via cholinergic modulation) (Tonini et al. 1989). Interestingly, *A. muciniphila* has been shown to be positively associated with gastrointestinal transit time (Gobert et al. 2016; Vandeputte et al. 2016). GABA could reach the CNS via blood stream as a lipophilic compound, being able to pass the blood brain barrier. Additionally, microbial GABA could affect the brain-gut axis by contributing the human GABA pools, especially as it has been shown that the microbiome can affect GABA receptor density in the CNS via the vagus nerve (Bravo et al. 2011). To establish whether and which role *A. muciniphila* and GABA may play a role in prodomal PD, further experimental studies will be required.

In order to move beyond mere cataloguing of microbial changes associated with diseases, pathway-based tools (Abubucker et al. 2012) have been developed, in which microbial sequences (or reads) are mapped, e.g., onto KEGG ontologies present in the KEGG database (Kanehisa et al. 2017). Using such tools, Bedarf et al reported decreased glucuronate degradation and an increase in tryptophan degradation and formate conversion (Bedarf et al. 2017). Similarly, Heinz-Buschart et al. reported 26 KEGG pathways to be altered in PD microbiomes (Heintz-Buschart et al. 2018). In our study, we complemented the compositional analysis with computational modelling to gain insight into potential functional, i.e., metabolic, consequences of changed microbe abundances in PD. The advantage of our approach is that the functional assignments may be more comprehensive than more canonical methods, such as KEGG ontologies because (1) the underlying genome-scale metabolic reconstructions have been assembled based on refined genome annotations and have been manually curated to ensure that the reaction and gene content is consistent with current knowledge about the microbe’s physiology, and (2) each of these reconstructions, alone or in combinations, are amenable to metabolic modelling and thus functional and metabolic consequences of a changed environment (e.g., nutrients or other microbes in the models) can be computed. These simulations are thus allowing to predict functional consequences and not only pathway or reaction enrichment, as typically done.

### Strength and limitation

Here, we present microbiome analyses in a large population-based, monocentric case-control study on PD from a defined area (Figure 5). Capitalising on the overall clinical spectrum of PD of the LuxPark cohort, which reflects a representative sample of PD patients of different disease stages from a defined geographical area, we demonstrated that microbial composition is not only altered in PD but also that the observed associations of PD with changes in the composition of the microbiome should be interpreted in the context of age, sex, BMI, and constipation. This information is of importance for clinical translation, highlighting the need for both, (i) a personalised and (ii) a holistic approach, to understand the role of microbial communities in PD pathogenesis. In a second step targeting the potential functional changes related to PD-associated microbiomes, we performed metabolic modelling based on the AGORA collection (Magnusdottir et al. 2017) of genome-scale metabolic reconstructions, allowing for the predictions of metabolite secretion profiles. Thus, our analyses facilitated a detailed investigation of the altered metabolism of PD-related microbial communities in the gut, pointing towards a role of the known pro-inflammatory species *B. wadsworthia* interacting with the host on sulphur metabolism. Hence, metabolic modelling provides a valuable tool for deciphering the metabolic activity of microbial communities in PD.

However, despite the partial confirmation of previous results by our study (Table 5), several limitations should be kept in mind. First, certain covariates were not investigated, such as diet, exercise, and smoking. Whether these covariates alter the PD-specific signature is yet to be analysed. Although our study belongs to the three largest studies performed yet on PD, our sample size was still too small to deliver insights on combinations of drugs. Furthermore, 16S RNA sequencing, as applied in our study, is not allowing analyses on the strain level and may lead to misclassifications (Janda and Abbott 2007), and follow-up studies based on shotgun sequencing are needed to further corroborate our results. However, our results are notably well aligned with a previous shotgun sequencing study (Bedarf et al. 2017), which would further support a role of 16S RNA sequencing as a cost-efficient screening method. Being cross-sectional in nature, causal inference is not possible. Consequently, although metabolic modelling has been numerous times been shown to correctly predict attributes of living systems (Oberhardt et al. 2009; Aurich and Thiele 2016; Nielsen 2017), our hypothesis on the role of *B. wadsworthia* in PD interlinking sulphur metabolism with disease severity requires experimental validation. To conclude, by combining metabolic modelling with comprehensive statistical analyses, we identified a promising research target in PD and refined the understanding of PD-related microbial changes.

## METHODS

### Description of the Luxembourg Parkinson’s study

For this study, data and biospecimen of the LuxPark cohort were utilised (Hipp et al. 2018). The Luxembourg Parkinson’s study includes a variegated group of patients with typical PD and atypical parkinsonism, and controls from Luxembourg and its neighbouring regions (Hipp et al. 2018). Controls were partly sampled among relatives of patients. The corresponding information on the family relation between controls and cases was not available. Cancer diagnosis with ongoing treatment, pregnancy, and secondary parkinsonism (drug-induced parkinsonism and parkinsonism in the frame of normotensive hydrocephalus) were exclusion criteria for enrolling in the patient or healthy control group. For 454 individuals (controls: n=248, PD: n=206) from the LuxPark cohort, stool samples were available and used for 16S RNA gene sequencing data (see below). Within LuxPark, controls were selected among spouses of chosen patients and volunteers and individuals from other independent Luxembourgish studies (Crichton and Alkerwi 2014; Ruiz-Castell et al. 2016). As we aimed to target specifically typical PD (IPD), we excluded all individuals with age below 50 (controls: n=47, PD: n=9) and all individuals with an unclear status of PD diagnosis or an atypical PD diagnosis (PD: n=47). PD patients were defined as typical PD, according to the inclusion criteria by the United Kingdom Parkinson’s Disease Society Brain Bank Clinical Diagnostic Criteria (Hughes et al. 1992). Furthermore, we excluded control patients with a United Parkinson’s Disease Rating Scale (UPDRS) III score above ten, except for one control where the high UPDRS III score was caused by an arm injury. Furthermore, we excluded control persons who took dopaminergic medications (n=5), and individuals who reported to have taken antibiotics in the last six months (controls: n=20, PD: n=13). Note that excluded observations behave sub-additive, because of overlap between the exclusion criteria (i.e. individuals below age 50 and taking antibiotics). Finally, 309 individuals (controls: n=162, cases: n=147) were included in the statistical analyses.

All study participants gave written informed consents, and the study was performed in accordance with the Declaration of Helsinki. The LuxPark study (Hipp et al. 2018) was approved by the National Ethics Board (CNER Ref: 201407/13) and Data Protection Committee (CNPD Ref: 446/2017).

### Measurements and neuropsychiatric testing

All patients and healthy controls were assessed by a neurologist, neuropsychologist or trained study nurse during the comprehensive battery of clinical assessment. Olfaction testing was conducted using the Sniffin’ Sticks 16-item version (SS) within the LuxPark cohort (Hipp et al. 2018). Antibiotics usage was defined as intake of antibiotic within the previous six months to stool collection. For assessing PD-related motor and non-motor symptoms, the UPDRS rating scales I-IV were used (Goetz et al. 2008). The severity of the disease was reflected by the Hoehn and Yahr staging (Hoehn and Yahr 1967). Non-motor symptoms were measured via the NMS questionnaire (Romenets et al. 2012). The use of medication was recorded, and PD-specific medication was classified into three classes, 1) levodopa, 2) dopamine receptor agonist, and 3) MAO-B/COMT inhibitors.

### Collection and processing of stool samples

All samples were processed following standard operating procedures (Lehmann et al. 2012; Mathay et al. 2015): stool samples were collected at home by patients using the OMNIgene.GUT stool tubes (DNA Genotek) and sent to the Integrated Biobank Luxembourg (IBBL) where one aliquot of 1 ml was used for DNA extraction. For the DNA extraction, a modified Chemagic DNA blood protocol was used with the MSM I instrument (PerkinElmer), the Chemagic Blood kit special 4 ml (Ref. CMG-1074) with a lysis buffer for faecal samples, and MSM I software. Samples were lysed using the SEB lysis buffer (included in the kit) and vortexed to obtain a homogenous suspension that was incubated for 10min at 70°C, then 5min at 95°C. Lysates (1.5mL) were centrifuged for five minutes at 10,000 g at RT. Supernatants were transferred to a 24XL deep-well plate. Plates were processed using the MSM I automated protocol.

### Analysis of the microbial composition with 16S rRNA gene sequencing

The V3-V4 regions of the 16S rRNA were sequenced at IBBL using an Illumina Platform (Illumina MiSeq) using 2×300bp paired-end reads (Hipp et al. 2018). The gene-specific primers targeted the V3 - V4 regions of the 16S rRNA gene. These primers were designed with Illumina overhang adapters and used to amplify templates from genomic DNA. Amplicons were generated, cleaned, indexed, and sequenced according to the Illumina-demonstrated 16S Metagenomic Sequencing Library Preparation Protocol with certain modifications. In brief, an initial PCR reaction contained at least 12.5 ng of DNA. A subsequent limited-cycle amplification step was performed to add multiplexing indices and Illumina sequencing adapters. Libraries were normalised, pooled, and sequenced on the Illumina MiSeq system using 2×300 bp paired-end reads.

The demultiplexed samples were processed merging forward and reverse reads and quality filtered using the dedicated pipeline “Merging and Filtering tool (MeFit)” (Parikh et al. 2016) with default parameters. To obtain a reliable microbial identification, identification to both genus and species taxonomic level was obtained using the SPINGO (SPecies level IdentificatioN of metaGenOmic amplicons) classifier (Allard et al. 2015) with default parameters. Relative abundances were computed, for each sample, using an R (R Foundation for Statistical Computing, Vienna, Austria) (Ihaka and Gentleman 1996) custom script. Briefly, for each sample, the counts of each genera/species were retrieved, and then the sum of the counts of all the genera/species was used to normalise to a total value of 1 each genera/species count.

### Personalised constraint-based modelling of microbial communities

AGORA consists of a set of 819 strains of microbes commonly found in the human gut (Magnusdottir et al. 2017; Noronha et al. 2019). To match species taxonomic resolution, we combined strain models of the same species in one species model (‘panSpeciesModel.m’) using the function ‘createPanModels.m’ of the microbiome modelling toolbox (Baldini et al. 2018). Briefly, reactions of multiple strains are combined into one pan-reconstruction. The pan-biomass reaction is built from the average of all strain-specific biomass reactions. Microbial abundances were mapped onto a set of 646 species performing an automatic name matching between SPINGO species taxonomic assignment and panSpecies names. A threshold for assessing the bacterial presence of a relative abundance value of 0.0001 was used to reduce the time of computations while limiting the order of magnitude simulations results of stoichiometric coefficients to ten. A total of 259 species overlapped between our set of species models and SPINGO species assignment when considering species identified at least in 10 % of samples (Supplementary Material). The retrieved microbial abundance information for each sample was integrated into a community modelling setup obtaining personalised microbiome models using the automated module of the microbiome modelling toolbox (Baldini et al. 2018) called mgPipe within the COBRA toolbox (Heirendt et al. 2019) (commit: b097185b641fc783fa6fea4900bdd303643a6a7e). Briefly, the metabolic models of the community members are connected by a common compartment, where each model can secrete/uptake metabolites. An average European diet was set as input for each microbiome model (Noronha et al. 2019). A community objective function was formulated based on the sum of each microbial model objective function and constrained to a lower bound of 0.4 per day and upper bound of one per day. A set of exchange reactions connects the shared compartment to the environment enabling to predict metabolite uptake and secretion flux rates (metabolic profiles/NMPCs) consistent with the applied constraints. The personalisation of each microbiome model was achieved by adjusting stoichiometric coefficients in the community biomass reactions to each sample’s relative microbial abundance and removing species undetected from the community models.

Relative reactions abundances were calculated by summing the number of species having the reaction in a microbiome model and scaling the sum by the respective species relative abundance. Community metabolic profiles of these microbial communities were assessed using flux variability analysis on the exchange reactions (Gudmundsson and Thiele 2010). AGORA microbial metabolic reconstructions used for the construction of the community models were downloaded from the VMH (www.vmh.life, (Noronha et al. 2019)). All computations were performed in MATLAB version 2018a (Mathworks, Inc.), using the IBM CPLEX (IBM, Inc.) solver through the Tomlab (Tomlab, Inc.) interface.

### Analyses of relative abundances

For descriptive statistics, metric variables were described by means and standard deviations, while nominal variables were described by proportions. Missing values were not imputed, and the pattern of missing values was not assessable via the ADA platform (Hipp et al. 2018). The read counts for each metagenomic feature (e.g., genera and species) were divided by total read counts such that relative abundances were retrieved. Relative abundances were checked for outliers. Observations with more than four standard deviations from the mean were excluded from analyses. Only genera and species detected in more than 50% of all samples were included in the analyses, resulting in 62 genera and 127 species.

The metagenomic data was analysed using fractional regressions as developed by (Papke and Wooldridge 1996). Fractional regressions, first applied to econometric problems, are semiparametric methods designed to model fractional data without the need of specifying the distribution of the response variable. Fractional regressions are further inherently robust against heteroscedasticity and can be parametrised in odds ratios, delivering convenient interpretations of the regression coefficients. All statistical models included technical covariates, batch, total read counts, and unclassified read counts (reads for which a taxonomic assignment was not possible independently from any threshold of confidence estimate value used). The read count variables were included into the statistical model, as it has been shown that normalisation by division can introduce bias if certain statistical assumptions implied by the application of division are not fulfilled (Hertel et al. 2018). In the case of metagenomic data, the effect of read counts would be removed by division if the observations would be sampled from a multinomial distribution. However, this is not a given as species and genera correlate amongst each other, violating the assumptions needed to construct multinomial distributions. In consequence, read count normalisation by division is prone to introduce a bias into metagenomic data; a potential bias, we corrected for by including the read counts as covariates into the model.

Before fitting the final statistical models, we explored the associations of basic covariates (age, sex, and BMI) with metagenomic features using fractional regressions as described above to avoid misspecifications of the statistical models. Since the data showed a high range in age and BMI, we checked for potential non-linear associations by including these variables into the models as restricted cubic splines (Harrell 2001) using three knots defined by the 5%-percentile, the median, and the 95%-percentile. As in the case for age, we found species with indications of non-linear age-associations with p<0.01, age was modelled in all analyses via restricted cubic splines.

All p-values are reported two-tailed. Statistical analyses were performed in STATA 14/MP (College Station, Texas, USA). Summary statistics of the performed analyses are given in the Supplementary files ‘Supplementary Tables’ 1-4.

### Differences between PD and controls in microbial composition and the influence of covariates

To analyse difference between genus abundances between PD and controls, fractional regressions were carried out with the relative abundance of the genus as the response variable, while including technical covariates, age (restricted cubic splines), sex, and BMI into the statistical modelling. The predictor of interest was the study group indicator variable. We corrected for multiple testing using the Benjamini-Hochberg procedure (Benjamini 2010) by setting the false discovery rate (FDR) to 0.05. Consequently, we corrected for 62 tests when reporting genera results. These analyses were repeated analogously for the taxonomic level of species, while correcting for multiple testing via the FDR.

Next, we explored the possibility of statistical interactions between basic covariates (age, sex, and BMI) and the group indicator. For these analyses, we once again modelled age and BMI via restricted cubic splines allowing for non-linear interaction terms. We only tested two-way interaction terms. All interaction terms were introduced simultaneously into the statistical model and tested on significance via a Wald test (Harrell 2001), correcting for multiple testing via the FDR. For the globally significant test, the single interaction terms were investigated to explore which covariate-group interaction contributed to the overall significance. For interpretation, the interaction terms were visually inspected by plotting the predictions conditional on technical covariates. These analyses were then rerun with species abundances as response variable instead of genus abundances.

We assessed the influence of constipation on the microbial composition. We introduced the binary predictor constipation (yes/no) as additional predictor into the model and the corresponding group-constipation interaction term. Both terms were tested simultaneously on zero with a Wald test. The analyses were once again adjusted for technical covariates, age (restricted cubic splines), sex, and BMI, and we corrected for multiple testing via the FDR.

### Analyses of within PD phenotypes in relation to microbial composition

We investigated the association pattern of medication and clinical features regarding the microbial composition. These analyses were only performed on the IPD cases, while controls were excluded from the analyses. First, we analysed the disease duration as measured in years between the date of the stool sampling and the year of the diagnosis. The analyses were conducted as before via fractional regressions with the genus abundances as the response variable, while adjusting for technical covariates, age (restricted cubic splines), sex, and BMI. Then, we assessed in separate analyses the UPDRS III score as an indicator for motor symptoms, the non-motor symptoms as measured by the NMS, the Hoehn-Yahr staging of the disease as a global measure of disease progression, and the sniff-score. All these analyses were performed adjusted for technical covariates, age (restricted cubic splines), sex, BMI, and disease duration. Each of these series of regression represents 62 test, which was accounted for using the FDR. The impact of medication was analysed by examining three classes of medication, a) levodopa, b) mono-amino oxidase/catechol-O-methyltransferase inhibitors, and c) dopamine receptor agonists. We generated three corresponding binary phenotypes (intake/no intake) and added these three variables simultaneously to the statistical model determining the significance of this add-on via a Wald test. We then tested each medication-class in separate analyses, strictly correcting for multiple testing via the FDR (186 tests in total). The analyses were performed adjusted for technical covariates, age (restricted cubic splines), sex, BMI, and disease duration.

### Statistical analyses of fluxes

The NMPCs were log transformed such that the skewness of the distribution was minimised (Box and Cox 1964)). This type of transformation was applied because of the very differently skewed distributions of the single NMPCs. Then, outliers were excluded using the 4-SD outlier rule as before. Only fluxes with more than 50% non-zero values were retained in analyses. Furthermore, NMPCs with distributions not suitable for statistical analyses (e.g., distributions with a high number of observations with exact the same numerical value) were excluded resulting in 129 NMPCs included into analyses.

The NMPCs were analysed with mixed linear regressions including the batch as random effects. Including the batch variable as a random effect has a higher statistical power in comparison to the fixed effect approach, but relies on more restrictive assumptions. We tested the corresponding random effect assumption by Hausman specification tests and found no indications of violations of the Hausman specification test. Note that this possibility to account for batch effects via random effects is not available with fractional regressions where batch effects were corrected via fixed effects.

We performed the same analyses as with the metagenomic data, with the sole exception of replacing the fractional regression model with the linear mixed model. In all other aspects, the analyses followed the same scheme.

### Analyses of species contribution to fluxes

To investigate the contribution of species and genera, we calculated for all included genera and all analysed fluxes the pairwise correlation and the corresponding variance contribution (the squared correlation). We classified every correlation above 0.5 (equal to 25% of variance contribution) as a strong correlation in accordance with classical classifications of effect size (Cohen 1988).

### Material availability

All 16S rRNA sequences can be requested from I.T. The mgPipe pipeline is available within the COBRA toolbox (https://github.com/opencobra/cobratoolbox), and the custom scripts with related documentation are available at the GitHub repository: https://github.com/ThieleLab/CodeBase/ND_collect.

## Acknowledgment

This study was funded by grants from the Luxembourg National Research Fund (FNR) within the National Centre of Excellence in Research (NCER) on Parkinson’s disease (FNR/NCER13/BM/11264123) and the PEARL programme (FNR/P13/6682797 to RK), by European Union’s Horizon 2020 research and innovation programme grant agreement No 692320 to RK, and by the European Research Council (ERC) under the European Union’s Horizon 2020 research and innovation programme (grant agreement No 757922) to IT.

